# Microglia in the developing retina couple phagocytosis with the progression of apoptosis via P2RY12 signaling

**DOI:** 10.1101/2020.01.30.925859

**Authors:** Zachary I. Blume, Jared M. Lambert, Anna G. Lovel, Diana M. Mitchell

**Affiliations:** Biological Sciences, University of Idaho, Moscow, ID 83844

**Keywords:** Microglia, phagocytosis, developmental apoptosis, retina, p2ry12, zebrafish

## Abstract

**Background:** Microglia colonize the developing vertebrate central nervous system coincident with detection of developmental apoptosis. Our understanding of apoptosis in intact tissue in relation to microglial clearance of dying cells is largely based on fixed samples, which is limiting given that microglia are highly motile and mobile phagocytes. Here, we used a system of microglial depletion and *in vivo* real-time imaging in zebrafish to directly address microglial phagocytosis of apoptotic cells during normal retinal development, the relative timing of phagocytosis in relation to apoptotic progression, and the contribution of P2RY12 signaling to this process.

**Results:** Depletion of microglia resulted in accumulation of numerous apoptotic cells in the retina. Real-time imaging revealed precise timing of microglial engulfment with the progression of apoptosis, and dynamic movement and displacement of engulfed apoptotic cells. Inhibition of P2RY12 signaling delayed microglial clearance of apoptotic cells.

**Conclusions:** Microglial engulfment of dying cells is coincident with apoptotic progression and requires P2RY12 signaling, indicating that microglial P2RY12 signaling is shared between development and injury response. Our work provides important *in vivo* insight into the dynamics of apoptotic cell clearance in the developing vertebrate retina and provides a basis to understand microglial phagocytic behavior in health and disease.

**Bullet Points:**

- Levels and location of developmental apoptosis in the zebrafish retina are elusive due to rapid and efficient clearance by microglia
- Microglial clearance of apoptotic cells is timed with the progression of apoptosis of the engulfed cell so that many cells are cleared in relatively early apoptotic stages
- P2RY12 signaling is involved in microglial sensing and clearance of cells undergoing normal developmental apoptosis, indicating shared signals in microglial responses to cell death in both healthy and injured tissue

**Grant Sponsors:** NIH NIGMS Grant No. P20 GM103408

## Introduction

Programmed cell death, a form of apoptosis that naturally occurs during development, is essential to proper tissue development, function, and homeostasis. During mammalian retinal development, programmed cell death occurs in large waves in a spatiotemporal fashion which is important to generate functional retinas ^1^. For example, in the postnatal mouse and rat retina, death of inner retinal neurons occurs over a period of several weeks, with the relative peak of cell death occurs at different timepoints for different inner retinal cell types ^2,3^. In zebrafish, comparably smaller waves have been observed and are thought to represent fine-tuning of developing retinal tissue, rather than elimination of a large number of developing neurons ^4^. In the embryonic mouse retina, detection of microglia ^5^, the resident phagocytic macrophage population, coincides temporally with detection of cell death ^1,6^. In the zebrafish, apoptotic cells can be detected in the brain and retina by 2 days post fertilization ^4,7–9^, which is also coincident with colonization of the central nervous system (CNS) by microglia ^7–12^. In addition, using the zebrafish model organism, recent studies indicate that normal developmental apoptosis is a significant factor driving macrophage entry into and establishment of microglia in the brain ^13,14^.

To date, clearance of apoptotic cells in the developing retina by microglia during normal retinal development is presumed based on studies from a variety of vertebrate organisms and based on fixed tissue samples ^4,12^. In addition, retinal Müller glia have been described to phagocytose dying cells during development and in response to tissue damage in several animal models^15–19^. A small body of work has emerged that uses the advantages of the zebrafish model organism to observe microglial phagocytosis and clearance of apoptotic neurons in the normally developing zebrafish brain ^20,21^; this work has focused mainly on microglial attachment to the apoptotic cell, phagosome formation/stabilization, and the subsequent intracellular events following engulfment by microglia rather than initial microglial sensing of the apoptotic cell or timing of engulfment. To our knowledge to date, there are no studies with real-time observations of clearance of apoptotic cells by microglia in the normally developing retina. Using fixed retinal tissue, work from Beihlmaier et al. ^4^ described a peak wave of apoptosis in the zebrafish retina around 3 days post fertilization (3 dpf), followed by a second wave around 7 dpf. Work from Herbomel et al. ^8^ indicates that colonization of the retina by microglia coincides temporally with the initial wave of developmental apoptosis. It is likely that rapid clearance of apoptotic cells in the developing retina makes numbers of apoptotic cells difficult to assess in fixed tissues, but such events have not been directly observed or documented, and the apparent levels of cell death in the developing zebrafish retina may be masked by microglial clearance.

A body of work has described signals released by apoptotic cells (“find-me” signals) that are sensed by and act as attractants for macrophage populations (reviewed in ^22,23^), though this is comparatively less well understood for microglia. Similarly, “eat me” signals have been described that facilitate microglial attachment to and phagocytosis of the apoptotic cell body ^22,24^. Overall, relatively little is known regarding the “find me” and “eat me” signals involved in normal developmental processes of the central nervous system. Further, it is not well described to what degree such signals are shared or distinct from those involved in responding to tissue damage, injury, or infection by microbes. For instance, the purinergic receptor P2RY12 (also called P2Y12) is expressed by microglia in mammals and zebrafish ^25–29^ and is involved in microglial responses to CNS injury through sensing ATP/ADP released by injured cells ^26,28^. In addition, purinergic signaling may be involved in colonization of the CNS by microglia ^13^, but a specific receptor was not identified. Indeed, a role for microglial P2RY12 in sensing cells undergoing normal developmental apoptosis, and thus leading to their clearance, is not well established.

Here, we use inducible depletion of microglia, *in vivo* real-time imaging, and pharmacological inhibition to show that microglia in the zebrafish retina rapidly sense, engulf, and clear cells undergoing developmental apoptosis, and that efficient microglial sensing of these apoptotic cells is dependent on P2RY12 signaling. We show that microglia in the developing zebrafish retina display exquisite timing of target cell engulfment relative to apoptotic progression, so that the majority of phagocytic events coincide with the progression of apoptosis. In addition, our live imaging work reveals dynamic movement of apoptotic cell bodies following microglial phagocytosis, as microglia continue active migration. Collectively, this work indicates that accurate levels of cell death are difficult to detect in the presence of active phagocytosis. In addition, using a potent and highly specific pharmacological inhibitor of P2RY12 signaling (PSB-0739 ^30,31^), we show that inhibition of P2RY12 signaling resulted in delayed microglial clearance of apoptotic cells in the retina. These findings suggest that similar to microglial responses to injury ^26,28^, P2RY12 signaling is also important in microglial clearance of cells undergoing programmed cell death during normal development. This work provides a new perspective on developmental apoptosis in the zebrafish retina and indicates that at least some “find me” signals released by dying cells are shared between injury and development.

## Results

### Depletion of microglia from developing zebrafish retinas results in accumulation of dying cells

Since phagocytic clearance of developmentally apoptotic cells during early retinal development is largely presumed, and to attempt to reveal more accurate levels of programmed cell death during early retinal development in zebrafish, we attempted to experimentally ablate microglial populations using a transgenic system. If microglial phagocytic clearance of apoptotic cells is indeed as presumed, such a manipulation would be expected to result in increased numbers of apoptotic cells because these cells would be expected to remain uncleared from the retina. Towards this goal, we utilized *mpeg1*:nfsBeYFP (also called *mpeg1*:NTReYFP) transgenic embryos ^30,31^ (Fig 1). This transgenic line carries a DNA construct in which bacterial nitroreductase (nfsB, also referred to as NTR) is driven by the *mpeg1* promoter. Therefore, *mpeg1*-expressing cells are sensitive to metronidazole-mediated ablation, because the metronidazole prodrug is converted to a toxic product by the nfsB enzyme. Similar systems (cell-type specific promoter driving bacterial NTR) have been used to ablate selected cell types in the zebrafish ^32–35^. Zebrafish microglia express *mpeg1*-driven transgenes ^11,15,31–33^, and we confirmed that microglia in retinas of *mpeg1*:nfsBeYFP zebrafish are YFP+ (Fig 1A, 1B), indicating that these microglial populations are susceptible to ablation by metronidazole (Mtz) treatment.

**Figure 1.**
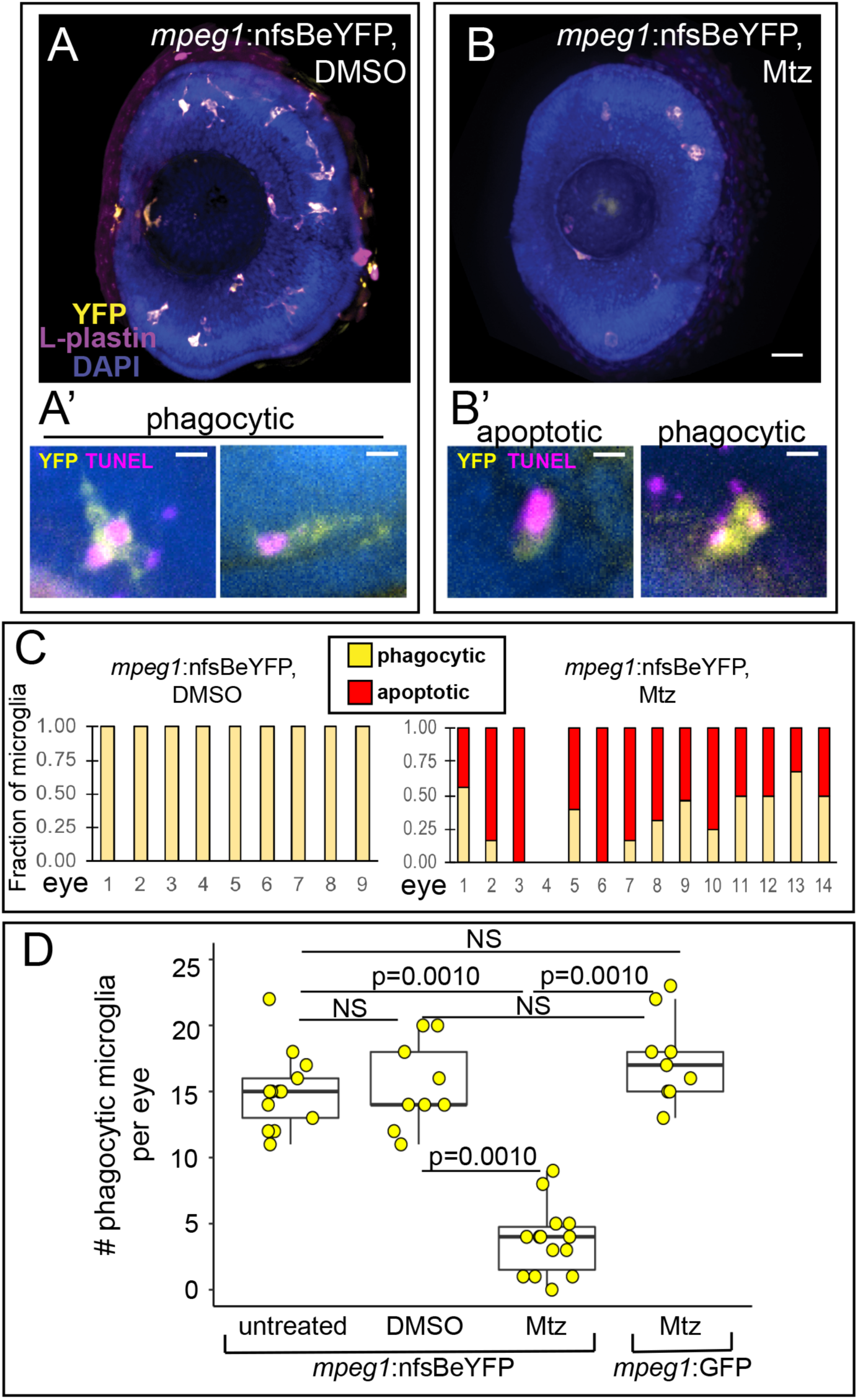
Mtz-mediated depletion of microglia in the retina of *mpeg1*:nfsBeYFP zebrafish embryos during development. A and B. Images of whole eyes from *mpeg1*:nfsBeYFP embryos following immersion in DMSO (A) or Mtz (B) from 24-72 hours post fertilization (hpf), stained for L-plastin (purple) and DAPI (blue) with YFP expression also visualized (yellow). All L-plastin+ cells in the developing retina express YFP (A), and these cells are reduced upon Mtz treatment (B). Scale bar in B = 20 microns and applies to A and B. A’. Examples of YFP+ microglia in control retinas with phagocytic morphology engulfing TUNEL+ dying cells (magenta). B’. Examples of morphology indicating apoptotic TUNEL+YFP+ microglia in retinas from *mpeg1*:nfsBeYFP retinas treated with Mtz (left) and remaining phagocytic YFP+ microglia engulfing TUNEL+ puncta (right). Scale bars in A’ and B’ = 5 microns. C. YFP+ microglia in *mpeg1*:nfsBeYFP retinas treated with DMSO (left) or Mtz (right) were assigned phagocytic or apoptotic morphology. Graphs show the fraction of microglia with each morphology in each eye examined. D. To determine the effective depletion, the number of phagocytic microglia in the retina in each indicated group was quantified. Box plots are shown for each group with each individual data point shown in yellow circles. A one-way ANOVA (p=2.04×10^−14^) was performed, followed by Tukey’s HSD post-hoc test. P values shown for pairwise comparison with p<0.05; NS=not significant. Phagocytic microglia were depleted only in *mpeg1*:nfs<u>B</u>eYFP embryos treated with Mtz. Samples sizes: *mpeg1*:nfsBeYFP untreated = 13 eyes from 13 fish, *mpeg1*:nfsBeYFP DMSO treated = 9 eyes from 5 fish, *mpeg1*:nfsBeYFP Mtz treated = 14 eyes from 10 fish, *mpeg1*:GFP Mtz treated = 9 eyes from 6 fish.

Since previous work has described a peak of developmental apoptosis in the zebrafish retina around 72 hours post fertilization (hpf) ^4^ and microglial colonization of the zebrafish retina beginning as early as 24 hpf ^8^, we decided to perform our experiments to deplete microglia during this initial peak of cell death and microglial colonization. To deplete microglia in the developing retina, embryos carrying the *mpeg1*:nfsBeYFP transgene were immersed in 10 mM Mtz from 24-72 hpf. Following this treatment, retinal microglia (which can be identified by co-label of *mpeg1-*driven reporter and the leukocyte marker L-plastin^3,5,34,35^) were reduced in number, but not completely ablated (compare Fig 1A and 1B). To better assess microglial depletion with this system, we analyzed the morphology of YFP+ microglia in combination with TUNEL staining and assigned each microglial cell as phagocytic or apoptotic (Fig 1A’ and 1B’). Phagocytic microglia were defined as YFP+ cells that displayed process extension(s) from the cell body which often showed phagocytic pouches, those with apparent vacuoles with or without TUNEL signal inside, those that appeared mobile, and/or those that did not have association with TUNEL+ signal. (Fig 1A’). Apoptotic microglia were defined as YFP+ cells with collapsed cytoplasm around TUNEL+ signal and a lack of vacuoles and extensions from the cell body. (Fig 1B’). All microglia in *mpeg1*:nfsBeYFP retinas treated with DMSO had phagocytic morphology, while the remaining microglia in *mpeg1*:nfsBeYFP retinas treated with Mtz showed both morphologies (Fig 1C). To determine the effective depletion of our system, and to ensure that Mtz did not kill microglia non-specifically, we determined the number of phagocytic microglia in retinas from *mpeg1*:nfsBeYFP fish (untreated, DMSO treated, or Mtz treated) and *mpeg1*:GFP fish (treated with Mtz) (Fig 1D). This analysis revealed that microglial numbers in *mpeg1*:nfsBeYFP retinas treated with Mtz were significantly reduced compared to the other groups (Fig 1D). It is worth noting that although significant, depletion varied in each sample (Fig 1D).

To examine accumulation of uncleared apoptotic cells in microglia-depleted retinas, we analyzed numbers of TUNEL+ cells in whole retinas (Fig 2). Increased total numbers of TUNEL+ cells were detected in microglia-depleted retinas (Fig 2A, 2A’, 2B, 2B’, 2C). Since TUNEL+ puncta were often found associated with microglia due to phagocytosis or microglia themselves that were dying in response to Mtz treatment (Fig 1A’, 1B’), we determined the number of TUNEL+ that were not associated with microglia and therefore likely representing uncleared dying cells. This analysis revealed that uncleared TUNEL+ cells were more numerous in microglia depleted retinas (Fig 2D), indicating that microglial clearance of dying cells in the developing retina likely masks the level of cell death that is detected at any single timepoint. Gross retinal structure did not display obvious qualitative changes when microglia were depleted as the cellular and plexiform layers of the developing retina were stereotypically structured (Fig 2E, 2F).

**Figure 2.**
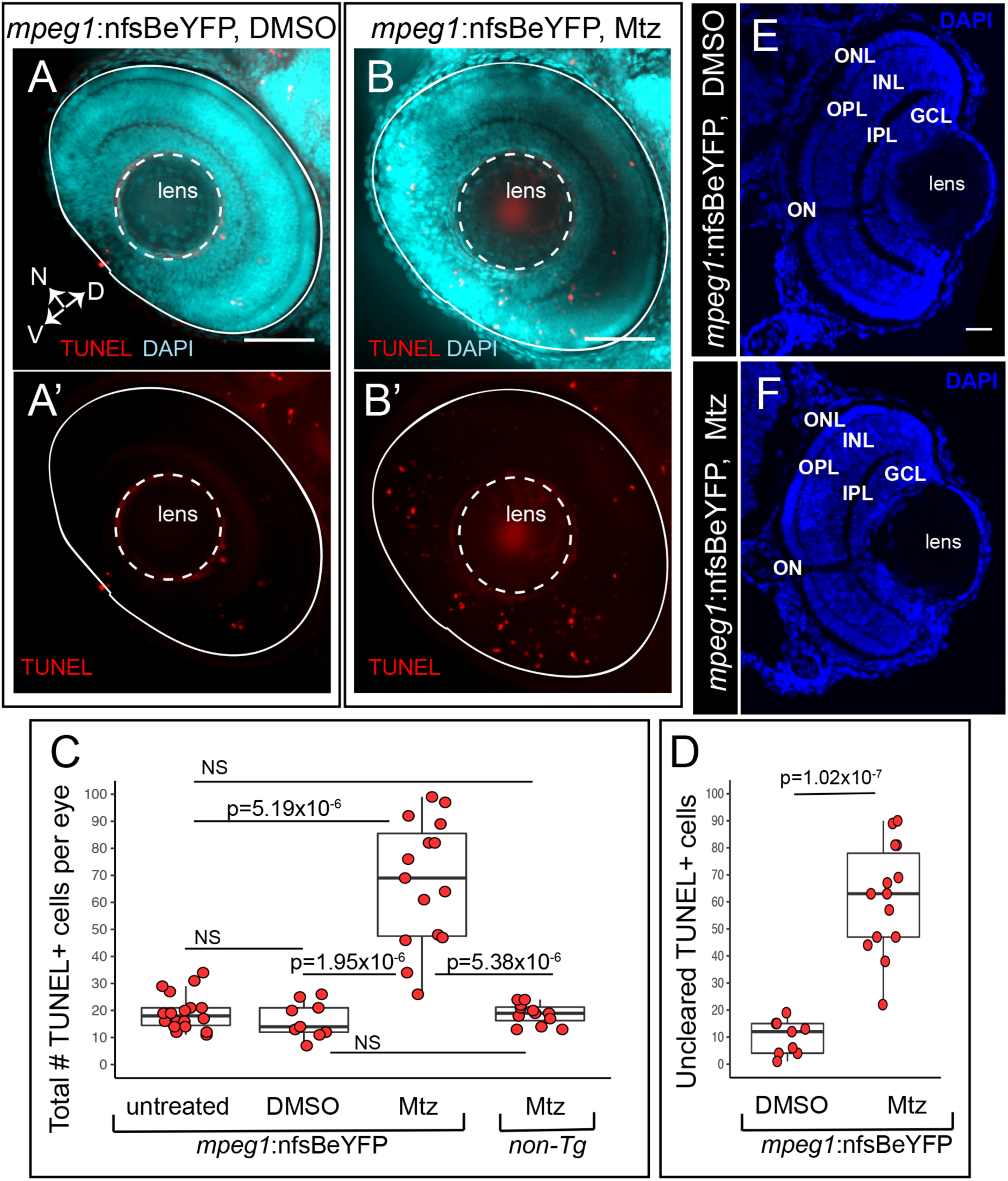
Accumulation of TUNEL+ cells in the retina when primitive microglia are depleted. Whole embryos were collected and processed for TUNEL (red) staining then DAPI (cyan) counterstaining. Images show selected z projections of whole eyes from *mpeg1*:nfsBeYFP embryos treated with DMSO (A, A’) or Mtz (B, B’). Orientation is indicated by the compass on bottom left of panel A, N=nasal, V=ventral, D=dorsal (applies to A-B’). The lens and boundary of the eye are indicated by white dashed outlines. C. The total number of TUNEL+ cells per eye was quantified for each indicated group. Box plots are shown for each group with each individual data point shown in red circles. A Welch’s one-way test (p=7.27×10^−7^) was performed, followed by Bonferroni post-hoc test. P values shown for pairwise comparison with p<0.05; NS=not significant. D. Since TUNEL signal is often associated with YFP+ microglia (see Fig 1A’ and B’), we determined the number of uncleared TUNEL+ cells in *mpeg1*:nfsBeYFP retinas treated with DMSO or Mtz by subtracting the number of microglia-associated TUNEL+ puncta from the total number of TUNEL+ cells. Box plots are shown for each group with each individual data point shown in red circles. P-value (Welch’s t-test) is shown. E and F. Comparison of gross retinal structure from retinal cryosections from control (D) or microglia depleted (E) eyes. DAPI (blue) was used to label cell nuclei and assess retinal structure. Stereotypical retinal structure is apparent for both control (E) and microglia-depleted retinas as evidenced by organized outer nuclear layer (ONL), inner nuclear layer (INL) and ganglion cell layer (GCL). The outer and inner plexiform layers (OPL and IPL, respectively) and location of the optic nerve (ON) are also apparent. Scale bar in E = 20 microns, applies to E and F. Samples sizes: *mpeg1*:nfsBeYFP untreated = 18 eyes from 15 fish, *mpeg1*:nfsBeYFP DMSO treated = 9 eyes from 5 fish, *mpeg1*:nfsBeYFP Mtz treated = 14 eyes from 10 fish, Non-Tg Mtz treated = 12 eyes from 6 fish.

It is worth noting that our analysis in Fig 1C suggests persistence of phagocytic microglia in *mpeg1*:nfsBeYFP retinas following Mtz treatment. It is possible that these remaining microglia may increase their phagocytic load to compensate for overall reduced numbers. Therefore, results from the *mpeg1*:nfsBeYFP system as utilized in our approach are difficult to fully interpret and further highlight the difficulties in interpreting levels of apoptosis in the presence of active phagocytosis. We therefore decided to use a live imaging approach to directly examine microglial phagocytosis in the developing retina.

### Rapid clearance of apoptotic cells by microglia in the developing zebrafish retina

To date, direct, real-time observation of microglial phagocytosis of cells undergoing developmental apoptosis in the retina is lacking. This is important to establish, as retinal Müller glia have been suggested to clear apoptotic cells in a variety of animals ^15–19^, while microglia in larval zebrafish eyes with induced, rod-specific cell death have been observed to engulf dying rods ^36^. In addition, the use of the zebrafish as a model for retinal degenerative disease is gaining popularity, for which an understanding of normal microglial phagocytic behaviors is required.

To directly observe clearance of apoptotic cells by retinal microglia in real-time, we used *in vivo* time-lapse imaging. Imaging of the developing zebrafish retina is facilitated by the fact that at the time of our analyses, the main volume of the eye consists nearly entirely of the lens and retina (Fig 3A). Embryos expressing the *mpeg1*:mCherry transgene were immersed in Acridine Orange (AO) solution beginning at approximately 54 hpf and were imaged live for 8 hours total (Fig 3, Movies 1 and 2). We chose this timing for imaging because it coincides with the initial peak of developmental apoptosis in the retina ^4^ and microglial colonization of the retina has begun ^8^. When complexed with DNA, AO displays green fluorescence and can be used to visualize apoptotic cells in whole tissue ^20,21,37–41^. In particular, AO has been used to image apoptotic cells in live zebrafish ^20,21,37,42^.

**Figure 3.**
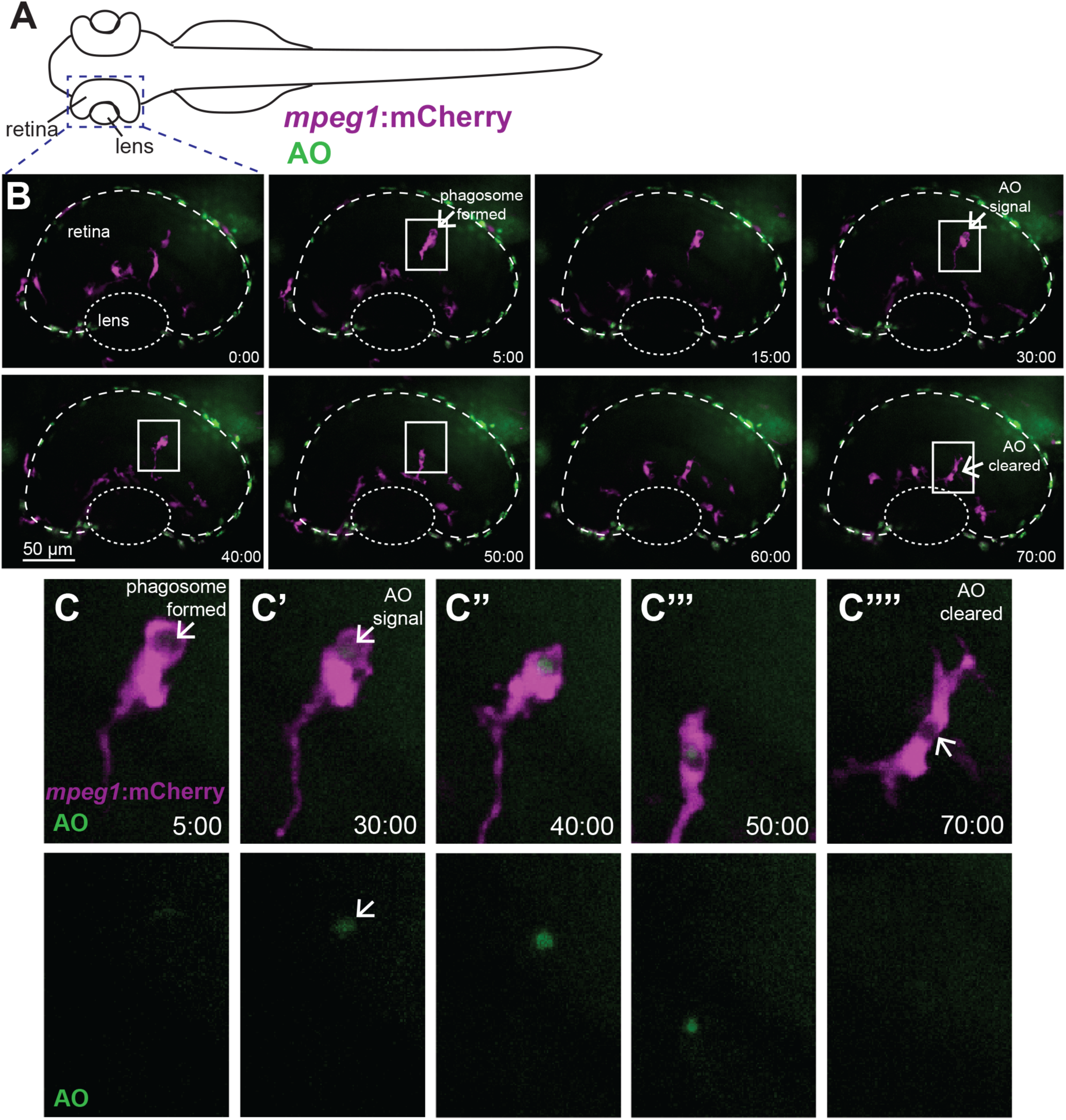
Real-time imaging and direct observation of apoptotic cell clearance by microglia during retinal development. A. Diagram of embryonic zebrafish as positioned for live imaging, indicating the developing eye/retina and region subjected to imaging. At the time of imaging, the developing zebrafish eye consists nearly entirely of the lens and retina. Retinas of *mpeg1*:mCherry embryos (to label microglia, magenta) beginning at approximately 54 hours post-fertilization (hpf) were imaged live using confocal microscopy. Acridine Orange (AO, green) was used to label apoptotic cells *in vivo*. Images were acquired every 5 minutes, with z stacks spanning the entire retina, for 8 hours total (ending at ∼62 hpf). B shows selected, flattened z projections and selected time-frames from a representative imaging session. Time stamps (min:sec) are shown at bottom right of each panel, with time 0:00 at the start of the selected frames. C-C’’’’ are enlarged insets of selected regions in B outlined by white boxes, showing *mpeg1*:mCherry and AO merged (top row) and AO alone (bottom row). In B and C, a phagosome is formed (arrow in second panel of B, arrow in C) ∼25 minutes prior to AO signal detection inside of the phagosome (arrow in fourth panel of B, arrow in C’ top and bottom). Clearance of the AO signal occurs approximately 40 minutes later (arrow, final panel of B and arrow in C’’’’). Initial AO signal detection was most frequently observed already associated with *mpeg1*:mCherry cells (see Fig 4C).

Microglia in the developing zebrafish retina were visualized using *mpeg1*:mCherry transgenic embryos (Movies 1 and 2, and Fig 3). Microglial phagosomes were visually apparent (Movie 2 and Fig 3B, 3C), and similar to Mazaheri et al.^20^, rapidly assembled phagosomes were observed to form compactly around the apoptotic cell (Movie 2 and Fig 3B, 3C). Over the 8 hours of imaging lasting approximately 54-62 hpf (n=6 embryos), we observed 21-29 AO+ cells per retina (Fig 4A). Essentially all of these AO+ cells were observed to be already engulfed by a microglial cell (Fig 4B). In fact, we most frequently observed that AO signal was initially detected while already associated with a microglial cell, often inside of a phagosome (Movies 1 and 2, Fig 3B, Fig 3C, and Fig 4C). Such observations indicate that engulfment of apoptotic cells by microglia occurs upon, or just prior to, late stages of the apoptotic process. Detection of AO signal in isolation (not already associated with a microglial cell) was only very rarely observed; such an event was only observed for one AO+ cell body in two of the six embryos imaged (Fig 4C). Degradation of engulfed apoptotic cells, as assessed by the time from AO signal detection (which was almost always already associated with a microglial cell) to disappearance, varied on an individual cell basis, with some degraded as rapidly as 10 minutes, and others as long as 120 minutes (Fig 4D).

**Figure 4.**
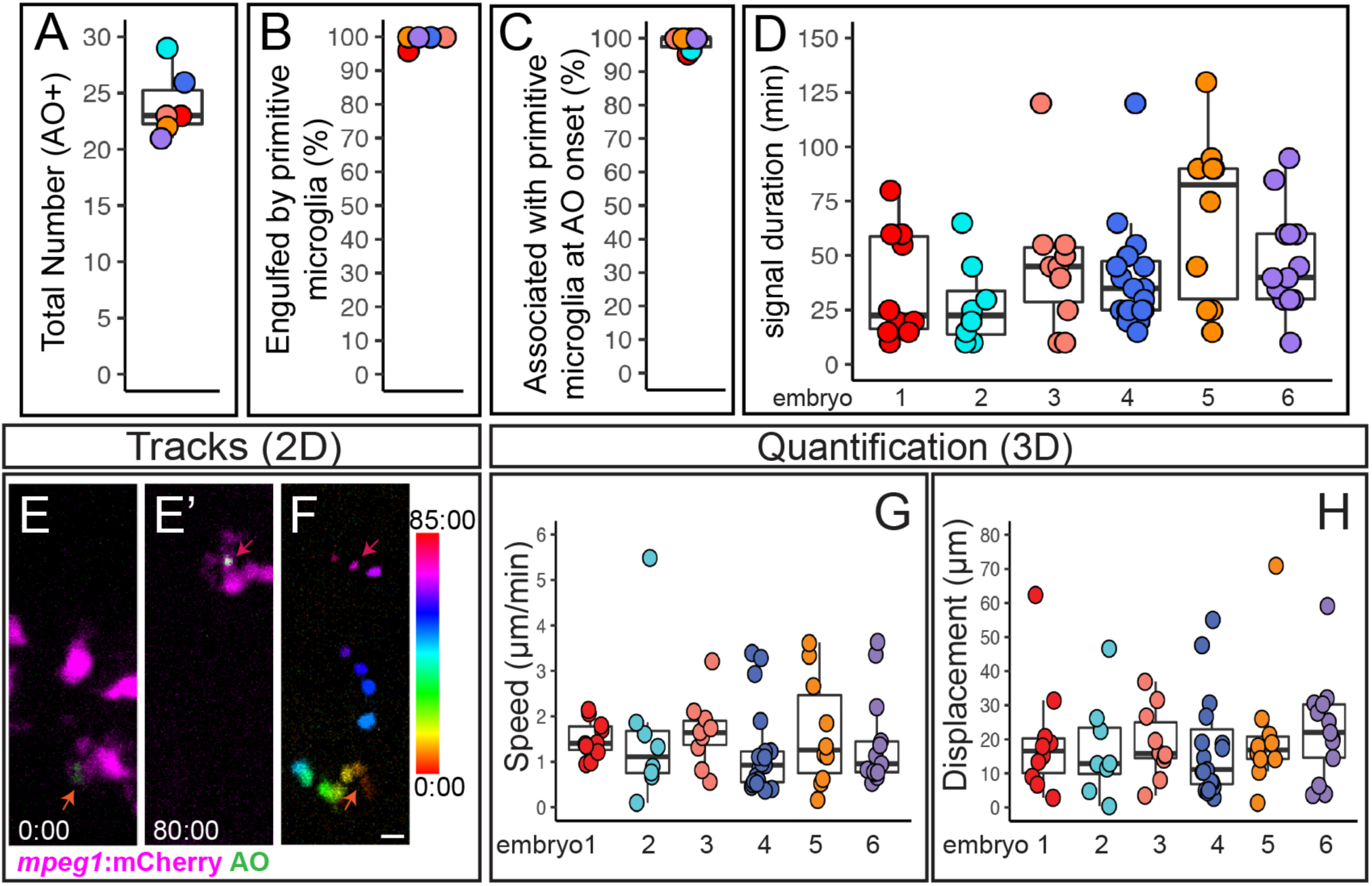
Quantification of cell death, microglial clearance, and apoptotic cell body movement detected by AO during retinal development. A-D. Each circle is color coded for each embryo imaged, and overlaid onto box plots. A. Total number of AO+ cells observed during live imaging from ∼54-62 hours post fertilization (hpf). B. The percent of AO+ cells that were observed to be engulfed by a microglial cell during the imaging session. C. For AO+ cells observed to be cleared by microglia, the percent showing association with a microglial cell at the onset of AO+ signal. D. Each circle represents individual cells from each embryo that was imaged. A selected number of individual AO+ cells, which were observed to appear and be cleared, during the imaging session were tracked in each embryo imaged. The duration of AO signal is shown for these individual AO+ cells. E-F represent max projections of selected, flattened z planes over a selected time-lapse. E. Shows the initial detection of an AO+ cell (green, indicated by red-orange arrow) associated with a microglial cell (microglia=magenta). E’. Shows the same AO+ cell (green, indicated by the magenta arrow) 80 minutes later, just before the AO signal is lost, after it had moved with the migrating microglial cell (magenta). F. The movement of the AO+ cell body was temporally color-coded to show its dynamic movement over time; the AO+ cell was associated with a microglial cell at all time frames. The color scale on the right indicates progression of time, where the first frame corresponds to 0:00 and the last frame is 85:00 (min:sec). Scale bar in bottom right of F = 5 microns and applies to E-F. G and H. To show a representation of dynamics of AO+ cell movement in association with microglia, a subset of AO+ cells were tracked in 3D using Imaris software. Quantifications show the speed (G) and total displacement (H) of individual AO+ cell bodies. Box plots are shown and circles are color coded to represent each embryo imaged. Each circle represents one AO+ cell that was tracked in each embryo that was imaged. Results are shown for n=6 different embryos.

During the 8 hours of live imaging, microglial cells in the retina were mobile and motile (Movies 1 and 2), continuing active migration following phagocytosis of target cells. Interestingly, engulfed AO+ cells were also observed to move dynamically while inside of microglial cells, as microglial processes were retracted and/or the microglia continued to migrate (Movie 1 and 2, Fig 3 and Fig 4E-H). We used 3D tracking in Imaris to follow a subset of the AO+ cells in the retina. We found that on average, engulfed AO+ cells moved at a speed of 1.5 micron/minute, which is consistent with previous reports of microglial migration and process extension/retraction ^20,43^. However, speeds varied on a per cell basis, with some moving as quickly as 3-5 microns/minute and some slower than 1 micron/minute (Fig 4G). Although recent work indicates that anesthesia may affect microglial motility in mice ^44,45^, it is not clear if this also has an effect in zebrafish. Nonetheless, movement of AO+ cells was only observed after microglial engulfment. We also determined the displacement of AO+ cells from initial signal until their disappearance (Fig 4H). Nearly all AO+ cells showed some measure of displacement, with some moving as far as 40-70 microns from their initial location (Fig 4H).

### Microglial sensing of apoptotic cells upon phosphatidyl serine (PtdSer) exposure

Since AO is limited to detecting cells in late stages of apoptosis, and given rapid clearance of apoptotic cells observed above, we imaged microglial engulfment in relation to exposure of phosphatidyl serine (PtdSer) on the outer membrane ^20,46,47^. PtdSer exposure on the cell membrane is an early apoptotic event ^48^. In the zebrafish CNS, PtdSer exposure occurs after Caspase activation, but prior to AO incorporation in the apoptotic pathway ^20^. In addition, PtdSer exposure is considered a potent “eat me” signal triggering microglial phagocytosis through attachment to a variety of cell surface receptors expressed on microglia ^22,49,50^. To label PtdSer+ cells *in vivo*, we used a transgenic zebrafish line, TBP-GAL4;UAS:secA5-YFP ^47^ (hereafter referred to as secA5-YFP). The secA5-YFP line carries two transgenic constructs. The TBP-GAL4 construct includes a TBP promoter (ubiquitously expressed) driving expression of GAL4, and GAL4 then binds to the UAS element (on the UAS:secA5-YFP construct) to drive expression of AnnexinA5 fused to YFP, resulting in ubiquitous expression of secreted AnnexinA5 fused to YFP ^47^. AnnexinV binds cooperatively to PtdSer ^51^ and has been used in a variety of techniques to label apoptotic cells (reviewed in ^52,53^). Thus, in secA5-YFP transgenic zebrafish, YFP signal is detected when the extracellular AnnexinA5-YFP binds to exposed PtdSer on the outer cell membrane of apoptotic cells. Similar to previous reports using and validating this transgenic line ^20,46,47,54^, fluorescent signal from the secA5 transgene product appeared to label most parts of the cell body and was detected around the presumed apoptotic cell body, as well as puncta on presumed neuronal processes (Movies 3 and 4, Fig 5A).

**Figure 5.**
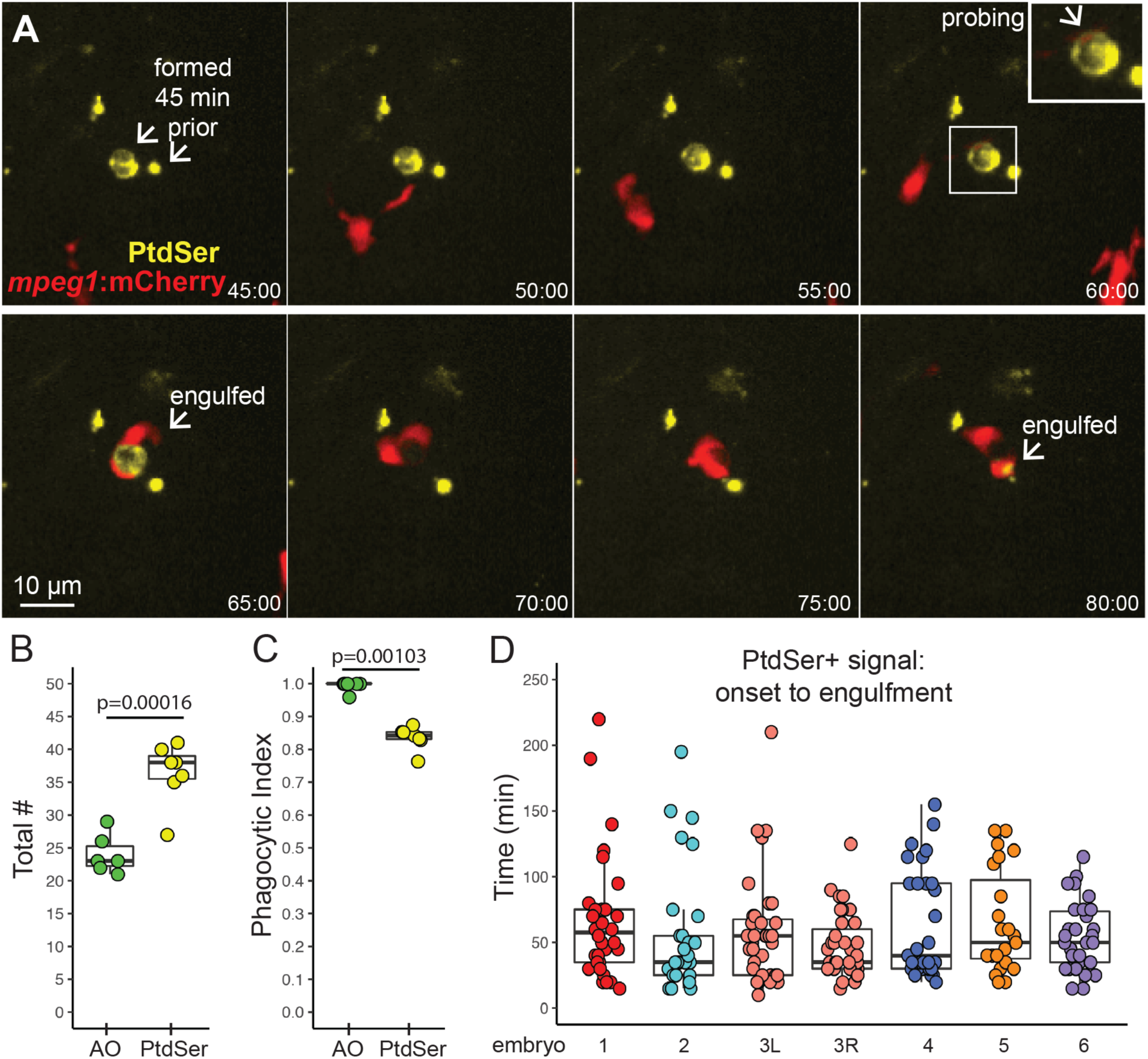
Microglial sensing of apoptotic neurons displaying Phosphatidyl Serine. A. Real-time imaging of microglia (red) sensing retinal neurons with exposed phosphatidyl serine (PtdSer, yellow). Selected panels are shown for an imaging session. Timestamp (hour:min, bottom right) is relative to YFP onset in the selection. In contrast to AO, YFP+ cell bodies were present (here, for ∼1 hour) prior to microglial sensing, probing, and engulfment. Scale bar applies to all images. Inset (upper right panel) shows enlargement of the PtdSer+ cell probed by a microglial process. B. Box plots show total number of PtdSer+ cells (n=6 embryos) over the imaging session, compared to AO+ numbers (n=6 embryos). Individual circles overlaid on box plots represent each sample. P value (Student’s 2-tailed t-test) is shown. C. Phagocytic index of apoptotic cells detected by AO or PtdSer. The phagocytic index is defined as the fraction of cells observed to be cleared by microglia during the imaging session. Box plots, with individual measurements overlaid (circles), are shown. P value from the generalized liner model with binomial family is shown. D. For each individual PtdSer+ cell counted, the time of PtdSer exposure from onset to microglial engulfment was calculated. Each box plot represents one eye from one embryo, with the exception of the salmon colored plots, which show two eyes from a single embryo. Each circle overlaid on the box plot represents one PtdSer+ cell.

Live imaging of *mpeg1*:mCherry;secA5-YFP double transgenic embryos from 54-62 hpf revealed that PtdSer exposure (as assessed by YFP signal) occurs prior to microglial engulfment; however, the duration of PtdSer presence prior to engulfment varies widely, from as little as 10 minutes to as long as 220 minutes (Movies 3 and 4, and Fig 5A and 5D). Notably, this is in contrast to AO signal, which as described above (Fig 3C, 4C) almost exclusively occurs already in association with a microglial phagosome. Typically, PtdSer signal disappeared soon after engulfment (Movies 3 and 4, and Fig 5A), ostensibly because the association of secA5-YFP and PtdSer was lost due to conditions in the microglial phagosome. Thus, we were not able to directly compare signal duration of PtdSer+ and AO+ cells, as these two signals differed temporally relative to microglial phagocytosis. In addition, microglia were observed to sometimes probe, retract, then later engulf PtdSer+ cells, and sometimes engulf presumed processes on later attempts (Movies 3 and 4, Fig 5A). Collectively, these observations confirm that, as in the brain ^20^, PtdSer exposure occurs before AO labeling of apoptotic cells in the retina. Of note, we also observed microglia “nibbling” at PtdSer+ cells (Movie 3), probably representing trogocytosis which has been recently described in microglial pruning of neuronal synapses ^55^.

Quantification of the total number of YFP+ (and therefore, PtdSer+) cell bodies over the 8-hour imaging session revealed higher numbers of PtdSer+ cells compared to those detected by AO in the same imaging period (Fig 5B). This suggests that many apoptotic cells are engulfed and cleared by microglia before apoptosis has progressed to relatively late stages. In agreement with this idea, the phagocytic index (i.e. the visualized association of microglia with apoptotic cells that are cleared during the imaging session) based on PtdSer signal is lower than that of AO signal (Fig 5C). Considering the dynamics of AO signal detection in relation to phagocytosis (Fig 3 and 4 above), these findings indicate that microglial engulfment of apoptotic cells is precisely timed to coincide with progression of the apoptotic pathway, so that apoptotic cells are engulfed just prior to or upon progression into late apoptosis.

### Inhibition of P2RY12 signaling delays microglial clearance of apoptotic cells in the developing zebrafish retina

The dynamics of apoptotic cell clearance as visualized by AO and PtdSer exposure indicate that a soluble signal is likely involved in microglial sensing of apoptosis progression, because the engulfment of the apoptotic cell is precisely timed to occur just prior to or to coincide with late apoptosis (Fig 3-5). Release of nucleotides by dying cells provides one class of “find me” signal that is sensed via purinergic receptors on phagocytes ^56^, but this is better characterized in systems of tissue injury or damage than in normally developing tissue. In particular, sensing of ATP/ADP released by damaged cells via binding to the P2RY12 receptor on microglia has been shown to be important for microglial responses to neuronal injury in the central nervous system in a variety of models ^26,28^. In addition, nucleotide signaling may be involved with colonization of microglia in the developing brain and retina downstream of developmental apoptosis^13^. We hypothesized that purinergic signaling through P2RY12 is also involved in sensing cells undergoing developmental apoptosis in the zebrafish retina. To test this idea, we used a highly potent, specific, and direct acting pharmacological inhibitor of the P2RY12 receptor, PSB-0739 ^28,30,31^. The zebrafish blood-brain barrier remains permeable by the time of our studies ^57^, as well as the retinal hyaloid vasculature ^58^. and until P2RY12 inhibition is applied embryos develop normally. Thus, immersion in PSB-0739 inhibitor is a feasible approach for inhibiting P2RY12 signaling during the time of interest.

Transgenic *mpeg1*:mCherry embryos were immersed in PSB-0739 for 1 hour prior to imaging, and live imaging was performed in the presence of PSB-0739 and AO (to detect apoptotic cells, Movies 5 and 6, Fig 6). In contrast to clearance observed in the retina of control animals (where AO+ signal was only twice detected in isolation, discussed above, Fig 4), inhibition of P2RY12 signaling resulted in the detection of several AO+ cells isolated from microglia (Movie 5, Fig 6A). We quantified and compared the total number of AO+ and PtdSer+ cells in retinas from control and PSB-0739 treated embryos (Fig 6B). PSB-0739 increased the number of late-stage apoptotic cells detected by AO, but did not significantly increase the number of PtdSer+ cells (early apoptosis) in the developing retina over the course of treatment and imaging (Fig 6B), indicating that P2RY12 inhibition prevented efficient apoptotic cell clearance but did not increase overall levels of cell death. To more directly address microglial clearance of apoptotic cells when P2RY12 signaling is inhibited, we next compared the phagocytic index of AO+ cells and PtdSer+ cells in control and PSB-0739 treated retinas (Fig 6C). Inhibition of P2RY12 signaling reduced the phagocytic index of AO+ cells to levels comparable to that of PtdSer+ cells, but the phagocytic index of PtdSer+ cells was not significantly affected by P2RY12 inhibition (Fig 6C). In addition, the duration of PtdSer signal from onset to engulfment was increased in the presence of PSB-0739 (Fig 6D). Collectively, these results indicate that we are observing apoptotic progression due to delayed, but not failed, clearance of apoptotic cells by microglia when P2RY12 signaling is inhibited. However, it is noteworthy that we occasionally observed instances, where in the presence of P2RY12 inhibition, PtdSer+ apoptotic cells were present for the entire 8-hour imaging session (Movie 6); this was never observed in control movies where the longest time-to-clearance of a PtdSer+ cell was approximately 220 minutes (∼3.7 hours). Since we did not observe these cells longer than 8 hours we cannot determine if their clearance failed or was substantially delayed. Collectively, our data indicates that P2RY12 signaling is required for rapid and efficient microglial sensing of apoptotic cells in the developing retina.

**Figure 6.**
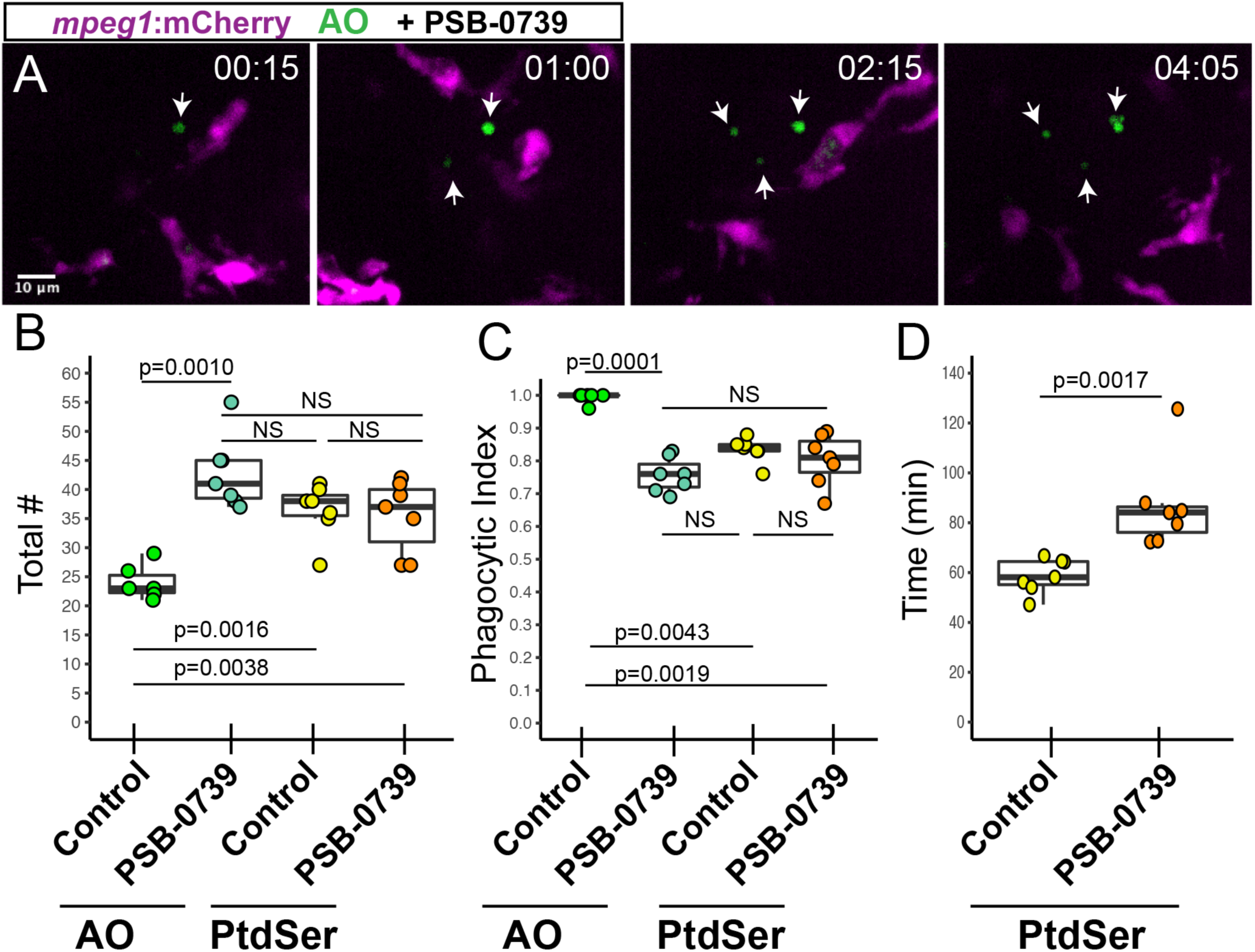
Pharmacological inhibition of P2RY12 signaling results in delayed clearance of apoptotic cells in the developing retina. *Mpeg1*:mCherry embryos were pretreated with PSB-0739 for 1 hour, then imaged with continual immersion in PSB-0739. Acridine Orange (AO, green) was used to label apoptotic cells *in vivo*. A. Selected z planes and timeframes are shown for imaging in the presence of PSB-0739. Isolated AO+ cells (arrows) are detected, which are ignored by microglia. Time stamp in upper right (hour:min) begins at appearance of the first AO+ cell (arrow, left panel). Scale bar in A applies to all panels. B. Box plots represent the total number of AO+ or PtdSer+ cells detected during 8 hours of imaging in control or PSB-0739 treated retinas (n=6-7 eyes from 6 embryos per group). Individual measurements are shown in circles and are overlaid on the box plots. A one-way ANOVA (p=0.00016) was performed, followed by Tukey’s HSD post-hoc test. P values shown for pairwise comparison with p<0.05; NS=not significant. An increased number of AO+ cells (late stage apoptosis) are detected in PSB-0739 treated retinas, but the total number of PtdSer+ cells detected in control and PSB-0739 treated retinas is not significantly different (NS). C. The phagocytic index (fraction of cells observed to be cleared by microglia) was determined for AO+ and PtdSer+ cells in control and PSB-0739 treated retinas. Box plots, with individual measurements (overlaid circles), are shown. A generalized linear model with binomial family was used, followed by one-way ANOVA (p=2.97×10^−12^) followed by Tukey’s post-hoc. P values are shown for pairwise comparisons with p<0.05, NS=not significant (n=6-7 eyes from 6 embryos per group). D. The time of PtdSer exposure from onset to engulfment was determined in control and PSB-0739 treated retinas. Box plots are shown, each overlaid circle represents the average of the measurements for each embryo. (n=7 eyes from 6 embryos per group) P value (Mann-Whitney U test) is shown. Results indicate that time from PtdSer exposure to microglial clearance is increased in the presence of PSB-0739.

## Discussion

The phagocytic potential of microglia has long been appreciated, and recent work has indicated such function is critical to CNS development ^20,21,59,60^. Our work provides direct *in vivo* observation of clearance of apoptotic cells by microglia in the normally developing zebrafish retina. In addition, we present validation of an inducible system for retinal microglia depletion with *mpeg1*:nfsBeYFP transgenic zebrafish (Fig 1). In our hands, this system results most consistently results in depletion, rather than ablation, of retinal microglial populations in embryonic fish. The reasons for incomplete ablation are unclear, but we speculate that this could be explained by several scenarios. Such scenarios include additional macrophage recruitment in response to induced and therefore increased cell death, the onset of *mpeg1*-expression in relation to the timing of our experiments resulting in a delay of effective levels of the NTR enzyme (nfsB), and/or that microglia are less sensitive to the Mtz-derived product than other cell types. In addition, it is possible that microglia that persist during the Mtz treatment may increase phagocytic activity to compensate for overall depleted microglial numbers. The inducible nature of this system remains advantageous for certain experimental goals, but the caveats should be well-considered.

The vertebrate retina is structurally and functionally conserved, with organization into stereotypical and clearly defined cellular and synaptic layers. Given the dynamic movement of apoptotic cells following microglial engulfment revealed by our work here (Fig 4E-H), with some cells displaced as many as 70 microns from their original location, assignment of the location of an apoptotic cell in fixed tissue may not be entirely accurate. By using two methods to detect apoptotic cells *in vivo* in relatively different stages of apoptosis, our data indicate that numbers of apoptotic cells present in the developing zebrafish retina are greater than previously appreciated from analyses of fixed tissue. Such findings provide a new perspective on programmed cell death in the developing zebrafish retina and support the zebrafish as a potential vertebrate model for such research.

Our work here indicates that microglia are the main cell type responsible for apoptotic cell clearance in the developing zebrafish retina, at least for the developmental timeframe analyzed, because we observe that essentially all of the late-stage (Acridine Orange+) cells and at least ∼80% of early-stage (PtdSer+) cells are engulfed by microglia (Fig 5C). Mazaheri et al. ^20^ found increased numbers of apoptotic cells in the developing zebrafish brain when microglia were depleted (using morpholino targeting of *pu.1*) and concluded that these increased numbers support primarily a clearance, rather than a survival, role for microglia in the developing CNS. In our work here, we show that in developing retinas with depleted microglial populations, there is an accumulation of TUNEL+ cells that likely also represent uncleared apoptotic cells (Fig 2), though due to the remaining microglia, is difficult to conclude based on the depletion system alone. Direct *in vivo* real-time imaging in our work confirms the microglial clearance function in the developing zebrafish retina (Figs 3-5). However, this does not exclude a possible phagocytic function for other cell types, such as Müller glia or possibly astrocytes in certain species ^61–63^, in the vertebrate retina. Indeed, Müller glial phagocytosis could occur at other developmental stages or in specific contexts of tissue damage or degeneration in zebrafish ^16–18^. Interestingly, a microglia-deficient zebrafish mutant has been described ^64^ which is viable to adult age. This gives rise to the questions of whether the presence of uncleared apoptotic cells in the zebrafish CNS is tolerated to some extent, or if other cell types (such as other glial populations) can assume this role in the absence of microglia. Such questions remain to be tested.

The apparent coupling of phagocytosis with apoptosis has been noted from samples of fixed tissues ^65–68^. It has been proposed that microglia promote progression of the apoptotic pathway within the target cell via a process called “phagoptosis” ^68–71^. Our findings confirm the coupling of phagocytosis and apoptosis *in vivo* in real time; however, similar to Mazaheri et al. ^20^, our data does not strongly support a role for microglial promotion of apoptosis within the target cell in the normally developing zebrafish CNS. Indeed, cells in late stages of apoptosis (detected by AO staining), but not early stages (detected by PtdSer), are increased in retinas where microglial sensing of apoptotic cells is inhibited (Fig 6), indicating that apoptosis progresses even when microglial engulfment is delayed. Our *in vivo* real-time imaging using these two different markers reveals that rather, microglial sensing and engulfment of apoptotic cells in the developing retina is exquisitely timed so that phagocytosis of target cells occurs upon, or even prior to, entry into late apoptosis.

In contrast to our observations in the retina (where we primarily observe apoptotic cell movement only after microglial engulfment), apoptotic cells in the developing neural tube themselves undergo directional movement to then arrive at phagocytic macrophages ^46^. Also noteworthy, a recent study found that microglia in the developing mouse retina phagocytose newborn retinal ganglion cells that were *not* undergoing programmed cell death ^60^. Such precise timing and diverse mechanisms for phagocytic clearance indicate that temporal coordination of signals released from, and presented by, both apoptotic and non-apoptotic cell targets and the evoked microglial attraction and phagocytic response differ in varying contexts and developmental timing. In addition, we also observed what appears to be microglial trogocytosis or “nibbling” at target cells (Movie 3), though the induction of and purpose of such activity in the current context remains to be determined.

In particular, our live imaging studies suggest that in order to time apoptotic engulfment precisely, a soluble mediator is involved in microglial localization of the apoptotic cell. We focused our attention on sensing of ATP/ADP through the P2RY12 receptor expressed on microglia ^25,26,28^, because nucleotide sensing via P2RY12 could provide a chemoattractant gradient allowing precise location of apoptotic cells in the developing retina. In addition, we were interested in determining if P2RY12 signaling, which is important in microglial response to CNS damage ^26,28,72^, is also an important mechanism in microglial sensing of cells undergoing developmental apoptosis. Indeed, in the presence of P2RY12 inhibition, clearance of apoptotic cells in the retina was delayed (Fig 6), indicating that P2RY12 signaling required for rapid and efficient microglia-mediated apoptotic cell clearance. Since clearance was delayed but not absent when P2RY12 signaling was inhibited, it is likely that, in addition to P2RY12 signaling, a variety of signals/receptors are important in microglial sensing of dying cells in the developing retina. In addition, it is possible that P2RY12 signaling is important in activation of microglial phagocytic machinery.

Rapid clearance of potentially detrimental material by microglia ensures proper CNS development and optimal function. A complete understanding of this dynamic process is crucial to understanding and mitigating pathology due to damage and degeneration of the CNS. It is also necessary in order to understand mechanisms involved with microglial sensing and phagocytosis of target cells in healthy tissue as opposed to disease. For example, dysregulated microglial phagocytosis may contribute to pathology in a variety of neurodegenerative diseases ^24,65,71,73–75^. In contrast, microglial phagocytosis limits secondary cell death in contexts of acute CNS injury ^76,77^ and is important in response to widespread retinal lesion in zebrafish ^78^. Therefore, a better understanding of molecular mechanisms and pathways functioning in microglial phagocytosis in healthy vs. damaged or diseased tissue are likely to yield important targets to modulate and balance microglial behavior *in vivo*. Our work has revealed, for the first time, *in vivo* dynamic processes of apoptotic cell clearance in the normally developing vertebrate retina and suggests that similar to microglial responses to trauma, this process requires purinergic signaling through the P2RY12 receptor. This work provides a basis for understanding microglial dynamics in healthy vs. damaged CNS tissue, and to further elucidate the mechanisms of target cell sensing and clearance.

### Experimental Procedures

#### Animals

All procedures using zebrafish were performed in compliance with IACUC (Institutional Animal Care and Use Committee) approved protocols at the University of Idaho. Adult zebrafish (Danio rerio) were maintained on a 14:10 light:dark cycle in 28.5°C recirculating, monitored system water, and were housed and propagated following Westerfield, 2007 ^79^. Embryos were collected into glass beakers in the morning, with light onset considered to be zero hours post fertilization (hpf), and water was refreshed daily until experimental endpoints. Zebrafish cannot be sexed before reproductive maturity; randomization and blinding were not used for the experiments.

Zebrafish lines used in this work include *mpeg1*:nfsBeYFP ^30^ (also called *mpeg1*:NTReYFP, obtained from Dr. Jefferey Mumm, Johns Hopkins University), *mpeg1*:GFP and *mpeg1*:mCherry ^32^ (obtained from Zebrafish International Resource Center, ZIRC), TBP-GAL4;UAS:secA5-YFP ^38,39^ (referred to as secA5-YFP throughout the manuscript, obtained from Dr. Randall Peterson, University of Utah), and a wild-type strain, referred to as SciH, originally obtained from Scientific Hatcheries (now Aquatica Tropicals) and maintained in-house.

#### Metronidazole treatments to deplete mpeg1+ cells

Metronidazole (Mtz, Acros Organics) solution was prepared fresh daily at 10 mM, 0.01% DMSO, dissolved in system water, immediately prior to treatment of embryos. Embryo clutches were collected from *mpeg1*:nfsBeYFP breeders or non-Mtz sensitive SciH line. Beginning at 24 hpf, the clutch of embryos from *mpeg1*:nfsBeYFP were split into two groups. After manual removal of chorions, one group was immersed in system water containing 10 mM Mtz, 0.01% DMSO, and the other group (control) was immersed in 0.01% DMSO. SciH (non-Mtz sensitive) were dechorionated and immersed in Mtz as a control for off-target effects of Mtz. Treatments were refreshed at 48 hpf, and embryos were collected at 72 hpf, for 48 hours total treatment. For those to be used in whole embryo immunofluorescent staining, 0.003% phenothiourea (PTU) was included in the water.

#### Whole embryo collection, processing, and immunofluorescent staining

For immunofluorescent staining, whole embryos were collected into 4% paraformaldehyde in PBS and fixed for one hour at room temperature, followed by three washes in PBST. Embryos were then transferred to blocking solution of 20% goat serum for 1 hour at room temperature with constant rotation, then to primary antibody (rabbit anti-L-plastin, 1:10,000, a kind gift of Dr. Michael Redd, University of Utah) and incubated overnight with constant rotation. Following three washes with PBST, embryos were then incubated in secondary antibody (Donkey anti-rabbit AlexaFluor647, 1:200) and DAPI (4.25 µM) for two hours with constant rotation, then washed two times in PBST followed by one wash in PBS. Prior to imaging, whole eyes were removed from the embryo head and mounted on a glass slide using Vectashield (Vector Labs), covered with a coverslip, and sealed with clear nailpolish.

#### Tissue collection and processing for retinal cryosections

Fixation and preparation of embryos for retinal cryosections were performed as previously described ^80^. Briefly, embryos were fixed in 4% paraformaldehyde containing phosphate buffered (pH=7.4) 5% sucrose for 1 hour at room temperature, then washed in phosphate buffered 5% sucrose, and transferred through a graded series ending in 20% sucrose solution. The following day, embryos were embedded in a 1:2 solution of OCT embedding medium (Sakura Finetek): 20% phosphate buffered sucrose, and frozen in isobutane supercooled with liquid N2. After freezing solid, tissue was sectioned at 10um thickness using a Leica CM3050 cryostat. Sections on slides were desiccated overnight, then stored at -20°C until stained with DAPI, then imaged.

#### TUNEL staining of whole embryos

For whole embryo TUNEL staining, embryos were collected into 4% paraformaldehyde in PBS containing 0.05% Tween and fixed overnight at 4°C with constant rocking. The following day, embryos were washed twice in PBS for 5 minutes each wash, then transferred to 50% methanol (5 minutes), then 100% methanol (5 minutes), then into fresh 100% methanol. Embryos were stored at -20C overnight then rehydrated the following day using the following graded series, for 5 minutes each at room temperature: 75% methanol:25% PBST, 50% methanol:50% PBST, 25% methanol:75% PBST, 100% PBST. After a second rinse in PBST, embryos were transferred to 0.1% sodium citrate solution for 15 minutes at room temperature with agitation. Proteinase K (supplier, final concentration) was then added and allowed to incubate for 5 minutes. Embryos were then washed in PBS containing 1% Triton-X100 for one hour, with the PBST refreshed every 20 minutes. During this time, reagents were prepared for TUNEL staining (Roche, TMR Red kit). 25 uL of enzyme solution was added to 225 ul of labeling solution in a microcentrifuge tube and mixed, then placed on ice. Embryos were removed from PBST wash and washed twice in PBS. Embryos were transferred to TUNEL reaction mixture in individual microcentrifuge tubes, with 2 embryos each placed in 20 uL of reaction mixture per tube. Tubes were incubated in the dark at 37C for 1.5 hours, then washed twice in PBS for 5 minutes each wash. Embryos were then stained with DAPI (4.25 µM) in PBST at room temperature for 20 minutes, then washed twice with PBS, and stored at 4C until imaging.

For imaging, embryos were transferred into a glass bottom culture dish (1.0 coverslip bottom, MatTek Corporation), PBS was removed, and embryos were immersed in glycerol. Embryos were oriented on their sides on the bottom of the dish’s coverslip.

#### In vivo real-time imaging

Embryos for live imaging were maintained in system water containing phenothiourea (PTU, 0.003%). At 48 hpf, embryos were screened for expression of fluorescent transgenes. Prior to treatment and live imaging, embryos were dechorionated and transferred to system water containing appropriate compound(s). In the case of acridine orange (AO) immersion, embryos were placed in system water containing AO (2 µg/mL, ThermoFisher) and PTU for 30 minutes prior to imaging. Immediately prior to live imaging, embryos were treated with tricaine to prevent movement (0.01% w/v final concentration), then transferred to a glass bottom culture dish (1.0 coverslip bottom, MatTek Corporation). Excess water was removed, and the embryos were embedded in 1.5% agarose and oriented with ventral side on the bottom of the dish. Once the agarose solidified, the dish was filled with fresh water containing AO, PTU, and tricaine. During live imaging, dishes were placed in a temperature-controlled chamber (Okolab, set to 28°C). The top and bottom of the eye was identified using DIC optics, and time-lapse images were acquired using 5 µm z stacks (z, ∼100 µm total z depth) obtained every 5 minutes (t) for 8 hours total. Viability of imaged embryos was confirmed following the imaging session by viewing the embryos under bright light and confirming that heartbeat and circulation appeared similar to that of unimaged embryos of the same age. Only data acquired from viable embryos was used for analyses.

#### Treatment with PSB-0739

PSB-0739 (Tocris) is a non-nucleotide competitive antagonist of the P2RY12 receptor ^30,31^ and is soluble in water. Following sorting for transgene expression and dechorionation as described above, half of the embryos were transferred to a new beaker containing PSB-0739 (50µM final concentration), Acridine Orange (2 µg/ml final concentration), and PTU (0.003%) in system water. The other half of the embryos (control) were transferred to a new beaker containing Acridine Orange (2 µg/ml final concentration), and PTU (0.003%) in system water. After 50 minutes, Tricaine was added to final concentration of 0.01% w/v to prevent movement of embryos prior to mounting for imaging. The embryos remained in the treatment solution with Tricaine for 10 minutes. Embryos were maintained at 28°C for the duration of treatment. When imaging with SecA5-YFP transgenics, treatments were prepared following the same procedure without the addition of Acridine Orange; the relevant volume was replaced with system water. Embryos were mounted for live imaging as described above and immersed in treatment solutions for the duration of imaging. Following the imaging session, viability of embryos was confirmed as above.

#### Microscopy, image acquisition, and image processing

Images were acquired using a Nikon Andor spinning disk confocal microscope equipped with a Zyla sCMOS camera and computer running Nikon Elements software. Imaging was performed using 20X (air) objective. For whole eyes, z stacks were obtained at 5 µm intervals. For live imaging, z stacks were obtained at 5 µm intervals. Time-lapse movies were visualized using Nikon Elements software, which was used to adjust brightness and contrast, annotate, then generate .avi files at a playback rate of 3 frames per second. The autofluorescent iridophores on the surface of the developing eye were used to determine the boundary of the retina for analysis and quantifications. Selected frames from timelapse movies were visualized using FIJI (ImageJ) or Nikon Elements Software. For imaging of 10 µm thick retinal cryosections, Z stacks were obtained at 2 µm intervals. For imaging of whole eyes, z stacks were obtained at 5 µm intervals. DIC optics were used to determine the boundaries of the eye on whole embryos. Images of retinal cryosections and whole eyes were visualized using FIJI (ImageJ) or Nikon Elements software.

#### Quantification and morphological scoring of YFP+ microglia

Images from whole eyes were processed using FIJI (ImageJ). The number of YFP+ microglia were counted in each individual z stack in the retina. Each YFP+ cell was assigned as phagocytic or apoptotic based on morphology. Phagocytic microglia were defined as YFP+ cells that displayed process extension(s) from the cell body which often showed phagocytic pouches, those with apparent vacuoles with or without TUNEL signal inside, those that appeared mobile, and/or those that did not have association with TUNEL+ signal. Apoptotic microglia were defined as YFP+ cells with collapsed cytoplasm around TUNEL+ signal and a lack of vacuoles and extensions from the cell body. To assess effective depletion, the number of phagocytic microglia (excluding apoptotic microglia) were compared across groups.

#### Quantification of TUNEL+ cells from whole eyes

Images from whole eyes imaged on intact embryos were processed using FIJI (ImageJ). The total number of TUNEL+ spots, considering coincidence with *mpeg1*:YFP or not, were counted in each individual z stack in the retina. The total number of TUNEL+ cells was determined, as well as the number of TUNEL+ spots associated with phagocytic or apoptotic microglia. The number of uncleared dying cells was determined by subtracting the number of TUNEL+ spots associated with microglia from the total number of TUNEL+.

#### Quantification of apoptotic cells from live imaging sessions

Quantifications were performed on regions of images within the developing retina. As discussed above, the presence of autofluorescent iridophores on the surface of the developing eye, as well as DIC images, were used to determine the boundary of the retina and lens in each separate z stack. Apoptotic cells, based on AO+ signal, were identified by puncta displaying fluorescence in the GFP+ channel. Apoptotic cells with exposed Phosphatidyl Serine (PtdSer, labeled by the SecA5-YFP transgene) were identified by YFP+ signal surrounding presumed cell bodies ^20,47^. A PtdSer+ cell was counted if the diameter of the YFP+ cell body was at least 4 microns. YFP+ puncta smaller than 4 microns were not considered cell bodies and therefore not counted. Using Nikon Elements software to view images, the total number of apoptotic cells appearing in the retina during a live imaging session were counted in all z stacks at all timepoints. The duration of AO+ or YFP+ signal was recorded for apoptotic cells that were observed to appear and disappear during the imaging session. Given that images were acquired every 5 minutes (described above), the time resolution for these quantifications is 5 minutes. To track AO+ cells in 3D, the Spot Tracker tool in Imaris (Bitplane) was used to determine AO+ cell movement (speed and displacement). Using the software’s automatic tracking, we were able to analyze a subset of AO+ cells in this manner.

#### Statistics

Box plots were chosen in order to show the distribution of data in each group. The box represents the first quartile (Q1, bottom line) to third quartile (Q3, top line) with the line inside of the box representing the median. The interquartile range (IQR) is determined by Q3-Q1. Whiskers extending above and below the box indicate Q3+1.5*IQR and Q1-1.5*IQR, respectively. Individual data points are shown overlaid on the box plots by colored circles.

Statistical analyses were performed using R coding environment. Prior to statistical comparison of data in which the total numbers of cells were quantified (i.e. microglia, TUNEL, AO, or PtdSer) or time was quantified (i.e. duration of PtdSer signal), each set of samples was evaluated for normality (using the Shapiro-Wilk test of normality) and equal variances (using Levene’s test for homogeneity of variances). For all statistical analyses, a p<0.05 was set as significance cut off. For pairwise comparisons, if both normality and homogeneity of variances was met for the samples to be compared, then a 2-tailed Student’s t-test was applied. If a comparison involved a set of samples that was not normally distributed, but variances were equal between the two groups, then the non-parametric Mann-Whitney U test was applied. If samples had normal distribution, but unequal variances, then a Welch’s t-test was used.

If comparisons were to be made across more than two samples, and both normality and homogeneity of variances were met for all groups, then a one-way ANOVA was used to indicate differences between means; Tukey’s post-hoc test was then applied for all pairwise comparisons. If groups showed unequal variances, then a Welch’s Heteroscedastic F Test was used as the one-way test with a Bonferonni post-hoc test for pairwise comparisons. For data involving proportions (i.e. phagocytic index), a generalized linear model with binomial family was used. For comparison of multiple groups, a one-way ANOVA was then used to indicate differences between means, with Tukey’s post-hoc test.

## Acknowledgements

Funding for this work was provided by an Idaho INBRE Developmental Research Project Grant under an Institutional Development Award (IDeA) from NIH NIGMS Grant No. P20 GM103408 awarded to DMM, start-up funds from the University of Idaho (DMM), and a University of Idaho Summer Undergraduate Research Fellowship (JML). We are grateful to Dr. Onesmo Balemba (Director of the UI Biological Sciences Opitical Imaging Core). We thank Dr. Jeff Mumm (Johns Hopkins University), Dr. Randall Peterson (University of Utah), and the Zebrafish International Resource Center (ZIRC) for providing transgenic zebrafish lines, and Dr. Michael Redd (University of Utah) for providing L-plastin antibody. We thank Dr. Deborah Stenkamp (University of Idaho) for providing critical review of a previous version of the manuscript, and Dr. Christopher Remien (University of Idaho) and Dr. Ben Ridenhour for assistance with generating box plots and statistical analyses.

## Movie Captions

**Movie 1.** *In vivo* real-time imaging of microglia (*mpeg1*:mCherry+, magenta) engulfing AO+ cells (green) in the developing zebrafish retina, 54 to 62 hours post fertilization. Overlap of magenta and green appears white. Timestamp shows hour:min and represents the entire imaging session. At this age, the developing eye consists nearly entirely of the retina and lens. The retina is outlined and region containing the lens is indicated. Autoflourescent iridiophores surround the surface of the developing eye which, in conjuction with DIC optics, establish the eye boundary.

**Movie 2.** Selected z and t frames of a microglial cell (magenta) engulfing an apoptotic cell labeled by AO. The phagosome forms prior to detection of AO signal (green) inside of the phagosome (arrow). Timestamp in upper left shows hour:min and represents the selected frames.

**Movie 3.** Selected z frames of microglia (red) engulfing cells with exposed Phosphatidyl Serine (PtdSer, yellow). Microglia engulf entire PtdSer+ cells (arrows), often probing the cells prior to engulfment. In addition, a microglia (red) nibbles bits of PtdSer off of a cell (yellow puncta), probably representing trogocytosis (upper left circle). Timestamp shows hour:min.

**Movie 4.** A microglia (red) engulfs a cell with exposed PtdSer (yellow). PtdSer is present for over an hour prior to microglial engulfment of the cell body, following microglial probing. A presumed cellular process is engulfed second; a smaller YFP+ puncta is left behind. Timestamp shows hour:min and represents the selected frames.

**Movie 5.** Microglia (magenta) ignore apoptotic cells (labeled by AO, green) in the presence of PSB-0739, a P2RY12 antagonist. Three apoptotic cells are detected (arrows) and are not engulfed by microglia, although some microglia have engulfed other AO+ cells as indicated by signal in microglial compartments. Timestamp in upper left shows hour:min.

**Movie 6.** Selected z frames showing a cell with surface PtdSer (yellow, arrow) that is transiently touched by a microglial cell (red), but never cleared, in the presence of PSB-0739, a P2RY12 antagonist. Over time, the YFP+ cell body appears to fragment. Timestamp in upper left shows hour:min and represents the entire imaging session.

## References

1. Péquignot M, Provost A, Sallé S, et al. Major role of BAX in apoptosis during retinal development and in establishment of a functional postnatal retina. Dev Dynam. 2003;228(2):231–238. doi:10.1002/dvdy.10376

2. Young RW. Cell death during differentiation of the retina in the mouse. J Comp Neurology. 1984;229(3):362–373. doi:10.1002/cne.902290307

3. Horsburgh GM, Sefton A. Cellular degeneration and synaptogenesis in the developing retina of the rat. J Comp Neurology. 1987;263(4):553–566. doi:10.1002/cne.902630407

4. Biehlmaier O, Neuhauss SC, Kohler K. Onset and time course of apoptosis in the developing zebrafish retina. Cell Tissue Res. 2001;306(2):199–207. doi:10.1007/s004410100447

5. ntos A, Calvente R, Tassi M, et al. Embryonic and postnatal development of microglial cells in the mouse retina. J Comp Neurology. 2007;506(2):224–239. doi:10.1002/cne.21538

6. Farah MH, Easter SS. Cell birth and death in the mouse retinal ganglion cell layer. J Comp Neurology. 2005;489(1):120–134. doi:10.1002/cne.20615

7. Rossi F, Casano A, Henke K, Richter K, Peri F. The SLC7A7 Transporter Identifies Microglial Precursors prior to Entry into the Brain. Cell Reports. 2015;11(7):1008–1017. doi:10.1016/j.celrep.2015.04.028

8. Herbomel P, Thisse B, Thisse C. Zebrafish Early Macrophages Colonize Cephalic Mesenchyme and Developing Brain, Retina, and Epidermis through a M-CSF Receptor-Dependent Invasive Process. Dev Biol. 2001;238(2):274–288. doi:10.1006/dbio.2001.0393

9. Bodeutsch N, Thanos S. Migration of phagocytotic cells and development of the murine intraretinal microglial network: An in vivo study using fluorescent dyes. Glia. 2000;32(1):91–101. doi:10.1002/1098-1136(200010)32:1<91::aid-glia90>3.0.co;2-x

10. Herbomel P, Thisse B, Thisse C. Ontogeny and behaviour of early macrophages in the zebrafish embryo. Dev Camb Engl. 1999;126(17):3735–3745.

11. 11. Martín-Estebané M, Navascués J, Sierra-Martín A, et al. Onset of microglial entry into developing quail retina coincides with increased expression of active caspase-3 and is mediated by extracellular ATP and UDP. Plos One. 2017;12(8):e0182450. doi:10.1371/journal.pone.0182450

12. Francisco-Morcillo J, Bejarano-Escobar R, Rodríguez-León J, Navascués J, Martín-Partido G. Ontogenetic Cell Death and Phagocytosis in the Visual System of Vertebrates. Dev Dynam. 2014;243(10):1203–1225. doi:10.1002/dvdy.24174

13. Casano A, Albert M, Peri F. Developmental Apoptosis Mediates Entry and Positioning of Microglia in the Zebrafish Brain. Cell Reports. 2016;16(4):897–906. doi:10.1016/j.celrep.2016.06.033

14. Xu J, Wang T, Wu Y, Jin W, Wen Z. Microglia Colonization of Developing Zebrafish Midbrain Is Promoted by Apoptotic Neuron and Lysophosphatidylcholine. Dev Cell. 2016;38(2):214–222. doi:10.1016/j.devcel.2016.06.018

15. Bejarano-Escobar R, Sánchez-Calderón H, Otero-Arenas J, Martín-Partido G, Francisco-Morcillo J. Müller glia and phagocytosis of cell debris in retinal tissue. J Anat. 2017;231(4):471–483. doi:10.1111/joa.12653

16. Sakami S, Imanishi Y, Palczewski K. Müller glia phagocytose dead photoreceptor cells in a mouse model of retinal degenerative disease. Faseb J. 2018;33(3):fj.201801662R. doi:10.1096/fj.201801662r

17. Bailey TJ, Fossum SL, Fimbel SM, Montgomery JE, Hyde DR. The inhibitor of phagocytosis, O-phospho-l-serine, suppresses Müller glia proliferation and cone cell regeneration in the light-damaged zebrafish retina. Exp Eye Res. 2010;91(5):601–612. doi:10.1016/j.exer.2010.07.017

18. Morris AC, Schroeter EH, Bilotta J, Wong RO, Fadool JM. Cone Survival Despite Rod Degeneration in XOPS-mCFP Transgenic Zebrafish. Invest Ophth Vis Sci. 2005;46(12):4762–4771. doi:10.1167/iovs.05-0797

19. Egensperger R, Maslim J, Bisti S, Holländer H, Stone J. Fate of DNA from retinal cells dying during development: uptake by microglia and macroglia (Müller cells). Dev Brain Res. 1996;97(1):1–8. doi:10.1016/s0165-3806(96)00119-8

20. Mazaheri F, Breus O, Durdu S, et al. Distinct roles for BAI1 and TIM-4 in the engulfment of dying neurons by microglia. Nat Commun. 2014;5(1):4046. doi:10.1038/ncomms5046

21. Peri F, Nüsslein-Volhard C. Live Imaging of Neuronal Degradation by Microglia Reveals a Role for v0-ATPase a1 in Phagosomal Fusion In Vivo. Cell. 2008;133(5):916–927. doi:10.1016/j.cell.2008.04.037

22. Elliott MR, Ravichandran KS. The Dynamics of Apoptotic Cell Clearance. Dev Cell. 2016;38(2):147–160. doi:10.1016/j.devcel.2016.06.029

23. Ravichandran KS. Find-me and eat-me signals in apoptotic cell clearance: progress and conundrums. J Exp Medicine. 2010;207(9):1807–1817. doi:10.1084/jem.20101157

24. Sierra A, Abiega O, Shahraz A, Neumann H. Janus-faced microglia: beneficial and detrimental consequences of microglial phagocytosis. Front Cell Neurosci. 2013;7:6. doi:10.3389/fncel.2013.00006

25. Butovsky O, Jedrychowski MP, Moore CS, et al. Identification of a unique TGF-β–dependent molecular and functional signature in microglia. Nat Neurosci. 2013;17(1):131–143. doi:10.1038/nn.3599

26. Haynes SE, Hollopeter G, Yang G, et al. The P2Y12 receptor regulates microglial activation by extracellular nucleotides. Nat Neurosci. 2006;9(12):nn1805. doi:10.1038/nn1805

27. Mitchell DM, Sun C, Hunter SS, New DD, Stenkamp DL. Regeneration associated transcriptional signature of retinal microglia and macrophages. Sci Rep-uk. 2019;9(1):4768. doi:10.1038/s41598-019-41298-8

28. Sieger D, Moritz C, Ziegenhals T, Prykhozhij S, Peri F. Long-Range Ca2+ Waves Transmit Brain-Damage Signals to Microglia. Dev Cell. 2012;22(6):1138–1148. doi:10.1016/j.devcel.2012.04.012

29. Oosterhof N, Holtman IR, Kuil LE, et al. Identification of a conserved and acute neurodegeneration-specific microglial transcriptome in the zebrafish. Glia. 2017;65(1):138–149. doi:10.1002/glia.23083

30. Baqi Y, Atzler K, Köse M, Glänzel M, Müller CE. High-affinity, non-nucleotide-derived competitive antagonists of platelet P2Y12 receptors. J Med Chem. 2009;52(12):3784–3793. doi:10.1021/jm9003297

31. Hoffmann K, Sixel U, Pasquale F, von Kügelgen I. Involvement of basic amino acid residues in transmembrane regions 6 and 7 in agonist and antagonist recognition of the human platelet P2Y12-receptor. Biochem Pharmacol. 2008;76(10):1201–1213. doi:10.1016/j.bcp.2008.08.029

32. White DT, Mumm JS. The nitroreductase system of inducible targeted ablation facilitates cell-specific regenerative studies in zebrafish. Methods San Diego Calif. 2013;62(3):232–240. doi:10.1016/j.ymeth.2013.03.017

33. Curado S, Stainier DY, Anderson RM. Nitroreductase-mediated cell/tissue ablation in zebrafish: a spatially and temporally controlled ablation method with applications in developmental and regeneration studies. Nat Protoc. 2008;3(6):948–954. doi:10.1038/nprot.2008.58

34. Curado S, Anderson RM, Jungblut B, Mumm J, Schroeter E, Stainier DY. Conditional targeted cell ablation in zebrafish: A new tool for regeneration studies. Dev Dynam. 2007;236(4):1025–1035. doi:10.1002/dvdy.21100

35. Pisharath H, Rhee JM, Swanson MA, Leach SD, Parsons MJ. Targeted ablation of beta cells in the embryonic zebrafish pancreas using E. coli nitroreductase. Mech Develop. 2007;124(3):218–229. doi:10.1016/j.mod.2006.11.005

36. White DT, Sengupta S, Saxena MT, et al. Immunomodulation-accelerated neuronal regeneration following selective rod photoreceptor cell ablation in the zebrafish retina. Proc National Acad Sci. 2017;114(18):E3719–E3728. doi:10.1073/pnas.1617721114

37. Weber T, Namikawa K, Winter B, et al. Caspase-mediated apoptosis induction in zebrafish cerebellar Purkinje neurons. Development. 2016;143(22):4279–4287. doi:10.1242/dev.122721

38. Hammerschmidt M, Pelegri F, Mullins M, et al. Mutations affecting morphogenesis during gastrulation and tail formation in the zebrafish, Danio rerio. Dev Camb Engl. 1996;123:143–151.

39. Abrams J, White K, Fessler L, Steller H. Programmed cell death during Drosophila embryogenesis. Dev Camb Engl. 1993;117(1):29–43.

40. Sidi S, Sanda T, Kennedy RD, et al. Chk1 suppresses a caspase-2 apoptotic response to DNA damage that bypasses p53, Bcl-2, and caspase-3. Cell. 2008;133(5):864–877. doi:10.1016/j.cell.2008.03.037

41. Furutani-Seiki M, Jiang Y, Brand M, et al. Neural degeneration mutants in the zebrafish, Danio rerio. Dev Camb Engl. 1996;123:229–239.

42. Paquet D, Bhat R, Sydow A, et al. A zebrafish model of tauopathy allows in vivo imaging of neuronal cell death and drug evaluation. J Clin Investigation. 2009;119(5):1382–1395. doi:10.1172/jci37537

43. Morsch M, Radford R, Lee A, et al. In vivo characterization of microglial engulfment of dying neurons in the zebrafish spinal cord. Front Cell Neurosci. 2015;9:321. doi:10.3389/fncel.2015.00321

44. Stowell RD, Sipe GO, Dawes RP, et al. Noradrenergic signaling in the wakeful state inhibits microglial surveillance and synaptic plasticity in the mouse visual cortex. Nat Neurosci. 2019;22(11):1782–1792. doi:10.1038/s41593-019-0514-0

45. Liu YU, Ying Y, Li Y, et al. Neuronal network activity controls microglial process surveillance in awake mice via norepinephrine signaling. Nat Neurosci. 2019;22(11):1771–1781. doi:10.1038/s41593-019-0511-3

46. van Ham TJ, Kokel D, Peterson RT. Apoptotic Cells Are Cleared by Directional Migration and elmo1-Dependent Macrophage Engulfment. Curr Biol. 2012;22(9):830–836. doi:10.1016/j.cub.2012.03.027

47. van Ham TJ, Mapes J, Kokel D, Peterson RT. Live imaging of apoptotic cells in zebrafish. Faseb J. 2010;24(11):4336–4342. doi:10.1096/fj.10-161018

48. Martin S, Reutelingsperger C, McGahon A, et al. Early redistribution of plasma membrane phosphatidylserine is a general feature of apoptosis regardless of the initiating stimulus: inhibition by overexpression of Bcl-2 and Abl. J Exp Medicine. 1995;182(5):1545–1556. doi:10.1084/jem.182.5.1545

49. Green D, Oguin T, Martinez J. The clearance of dying cells: table for two. Cell Death Differ. 2016;23(6):915–926. doi:10.1038/cdd.2015.172

50. Barth ND, Marwick JA, Vendrell M, Rossi AG, Dransfield I. The “Phagocytic Synapse” and Clearance of Apoptotic Cells. Front Immunol. 2017;8:1708. doi:10.3389/fimmu.2017.01708

51. Janko C, Jeremic I, Biermann M, et al. Cooperative binding of Annexin A5 to phosphatidylserine on apoptotic cell membranes. Phys Biol. 2013;10(6):065006. doi:10.1088/1478-3975/10/6/065006

52. Brumatti G, Sheridan C, Martin SJ. Expression and purification of recombinant annexin V for the detection of membrane alterations on apoptotic cells. Methods San Diego Calif. 2008;44(3):235–240. doi:10.1016/j.ymeth.2007.11.010

53. Logue SE, Elgendy M, Martin SJ. Expression, purification and use of recombinant annexin V for the detection of apoptotic cells. Nat Protoc. 2009;4(9):1383–1395. doi:10.1038/nprot.2009.143

54. Svahn AJ, Graeber MB, Ellett F, et al. Development of ramified microglia from early macrophages in the zebrafish optic tectum. Dev Neurobiol. 2013;73(1):60–71. doi:10.1002/dneu.22039

55. Weinhard L, di Bartolomei G, Bolasco G, et al. Microglia remodel synapses by presynaptic trogocytosis and spine head filopodia induction. Nat Commun. 2018;9(1):1228. doi:10.1038/s41467-018-03566-5

56. Elliott MR, Chekeni FB, Trampont PC, et al. Nucleotides released by apoptotic cells act as a find-me signal to promote phagocytic clearance. Nature. 2009;461(7261):282. doi:10.1038/nature08296

57. Fleming A, Diekmann H, Goldsmith P. Functional Characterisation of the Maturation of the Blood-Brain Barrier in Larval Zebrafish. Plos One. 2013;8(10):e77548. doi:10.1371/journal.pone.0077548

58. Hartsock A, Lee C, Arnold V, Gross JM. In vivo analysis of hyaloid vasculature morphogenesis in zebrafish: A role for the lens in maturation and maintenance of the hyaloid. Dev Biol. 2014;394(2):327–339. doi:10.1016/j.ydbio.2014.07.024

59. Schafer DP, Lehrman EK, Kautzman AG, et al. Microglia Sculpt Postnatal Neural Circuits in an Activity and Complement-Dependent Manner. Neuron. 2012;74(4):691–705. doi:10.1016/j.neuron.2012.03.026

60. Anderson SR, Zhang J, Steele MR, et al. Complement Targets Newborn Retinal Ganglion Cells for Phagocytic Elimination by Microglia. J Neurosci Official J Soc Neurosci. 2019;39(11):2025–2040. doi:10.1523/jneurosci.1854-18.2018

61. Watabe K, Osborne D, Kim SU. Phagocytic Activity of Human Adult Astrocytes and Oligodendrocytes in Culture. J Neuropathology Exp Neurology. 1989;48(5):499–506. doi:10.1097/00005072-198909000-00001

62. Wakida NM, Cruz GS, Ro CC, et al. Phagocytic response of astrocytes to damaged neighboring cells. Plos One. 2018;13(4):e0196153. doi:10.1371/journal.pone.0196153

63. Morizawa YM, Hirayama Y, Ohno N, et al. Reactive astrocytes function as phagocytes after brain ischemia via ABCA1-mediated pathway. Nat Commun. 2017;8(1):28. doi:10.1038/s41467-017-00037-1

64. Shiau CE, Kaufman Z, Meireles AM, Talbot WS. Differential Requirement for irf8 in Formation of Embryonic and Adult Macrophages in Zebrafish. Plos One. 2015;10(1):e0117513. doi:10.1371/journal.pone.0117513

65. Fricker M, Neher JJ, Zhao J-W, Théry C, Tolkovsky AM, Brown GC. MFG-E8 Mediates Primary Phagocytosis of Viable Neurons during Neuroinflammation. J Neurosci. 2012;32(8):2657–2666. doi:10.1523/jneurosci.4837-11.2012

66. Sierra A, Encinas JM, Deudero J, et al. Microglia Shape Adult Hippocampal Neurogenesis through Apoptosis-Coupled Phagocytosis. Cell Stem Cell. 2010;7(4):483–495. doi:10.1016/j.stem.2010.08.014

67. Neher JJ, Neniskyte U, Zhao J-W, Bal-Price A, Tolkovsky AM, Brown GC. Inhibition of Microglial Phagocytosis Is Sufficient To Prevent Inflammatory Neuronal Death. J Immunol. 2011;186(8):4973–4983. doi:10.4049/jimmunol.1003600

68. Marín-Teva J, Dusart I, Colin C, Gervais A, van Rooijen N, Mallat M. Microglia Promote the Death of Developing Purkinje Cells. Neuron. 2004;41(4):535–547. doi:10.1016/s0896-6273(04)00069-8

69. Brown GC, Neher JJ. Microglial phagocytosis of live neurons. Nat Rev Neurosci. 2014;15(4):209–216. doi:10.1038/nrn3710

70. Vilalta A, Brown GC. Neurophagy, the phagocytosis of live neurons and synapses by glia, contributes to brain development and disease. Febs J. 2018;285(19):3566–3575. doi:10.1111/febs.14323

71. Abiega O, Beccari S, Diaz-Aparicio I, et al. Neuronal Hyperactivity Disturbs ATP Microgradients, Impairs Microglial Motility, and Reduces Phagocytic Receptor Expression Triggering Apoptosis/Microglial Phagocytosis Uncoupling. Plos Biol. 2016;14(5):e1002466. doi:10.1371/journal.pbio.1002466

72. Lou N, Takano T, Pei Y, Xavier AL, Goldman SA, Nedergaard M. Purinergic receptor P2RY12-dependent microglial closure of the injured blood–brain barrier. Proc National Acad Sci. 2016;113(4):1074–1079. doi:10.1073/pnas.1520398113

73. Zhao L, Zabel MK, Wang X, et al. Microglial phagocytosis of living photoreceptors contributes to inherited retinal degeneration. Embo Mol Med. 2015;7(9):1179–1197. doi:10.15252/emmm.201505298

74. Neumann H, Kotter M, Franklin R. Debris clearance by microglia: an essential link between degeneration and regeneration. Brain. 2009;132(2):288–295. doi:10.1093/brain/awn109

75. Diaz-Aparicio I, Beccari S, Abiega O, Sierra A. Clearing the corpses: regulatory mechanisms, novel tools, and therapeutic potential of harnessing microglial phagocytosis in the diseased brain. Neural Regen Res. 2016;11(10):1533–1539. doi:10.4103/1673-5374.193220

76. Herzog C, Garcia L, Keatinge M, et al. Rapid clearance of cellular debris by microglia limits secondary neuronal cell death after brain injury in vivo. Development. 2019;146(9):dev174698. doi:10.1242/dev.174698

77. Okunuki Y, Mukai R, Pearsall EA, et al. Microglia inhibit photoreceptor cell death and regulate immune cell infiltration in response to retinal detachment. Proc National Acad Sci. 2018;115(27):201719601. doi:10.1073/pnas.1719601115

78. Mitchell DM, Lovel AG, Stenkamp DL. Dynamic changes in microglial and macrophage characteristics during degeneration and regeneration of the zebrafish retina. J Neuroinflamm. 2018;15(1):163. doi:10.1186/s12974-018-1185-6

79. Westerfield M. The zebrafish book. A guide for the laboratory use of zebrafish (Danio rerio). 2007.

80. Mitchell DM, Stevens CB, Frey RA, et al. Retinoic Acid Signaling Regulates Differential Expression of the Tandemly-Duplicated Long Wavelength-Sensitive Cone Opsin Genes in Zebrafish. Plos Genet. 2015;11(8):e1005483. doi:10.1371/journal.pgen.1005483

